# Mbd3 and deterministic reprogramming

**DOI:** 10.1101/013904

**Authors:** Paul Bertone, Brian Hendrich, José C.R. Silva

**Author notes:** Correspondence (PB), (BH), (JCRS).

## Abstract

Embryonic development requires the activity of the Nucleosome Remodeling and Deacetylase (NuRD) complex. NuRD functionality can be ablated by rendering cells devoid of methyl-CpG-binding domain protein 3 (Mbd3), a critical component that confers stability to the complex^1^. Previous studies noted that *Mbd3*-/- embryonic stem (ES) cells misregulate a subset of pluripotency-associated genes, and subsequently fail to engage in cell differentiation into embryonic lineages when self-renewal requisites (e.g. LIF) are withdrawn from culture media^2–3^. Components of the NuRD complex have been shown to interact with Oct4 and Nanog, two important transcription factors operative in the production of iPS cells^4–7^. Thus, elucidating the role of Mbd3/NuRD in the reprogramming process is of relevance to the field.

Rais *et al.* reported the remarkable observation that suppressing formation of the NuRD complex by depleting Mbd3 promotes near 100% induction of cells reprogrammed to pluripotency^8^. This was an important result, as typical iPS cell conversion rates are extremely low; such an increase in reprogramming efficiency would represent a considerable advance in the production of iPS cells for research and therapeutic applications. However, concurrent and independent work from our labs obtained contrasting results, where a profound reduction in reprogramming efficiency was observed from cells where *Mbd3* had been ablated^9^.

We sought to understand this discrepancy, in part by analyzing the data provided by Rais *et al*. The study employs cells containing a single functional allele of *Mbd3* that can be conditionally deleted (*Mbd3*^fl/-^)^10^, and that are also transgenic for several constructs inserted into the genome by random integration: a doxycycline (DOX)-inducible polycistronic reprogramming cassette, a promoter-driven Oct4-GFP reporter, and a constitutive mCherry reporter used for quantification of colony sizes and single-cell deposition by flow sorting. Performance of *Mbd3*^fl/-^ cells in reprogramming assays was described in Rais *et al*. relative to *Mbd3*^+/+^ counterparts.

A gene expression dataset was produced for the study, where *Mbd3*^fl/-^ and *Mbd3*^+/+^ mouse embryonic fibroblasts (MEFs) were profiled on Affymetrix arrays during a reprogramming timecourse. Comprehensive analysis of this experiment is precluded by its design: only four time points are represented, one of those is inconsistent between experiment and control samples (taken at 8 vs 11 days), and no replicates were provided. Nevertheless, differential expression analysis reveals highly similar outcomes in each condition (Fig. 1A), with little divergence among pluripotency genes during induction of *Mbd3* heterozygous and wild-type cells (Fig. 1B).

**Figure 1.**
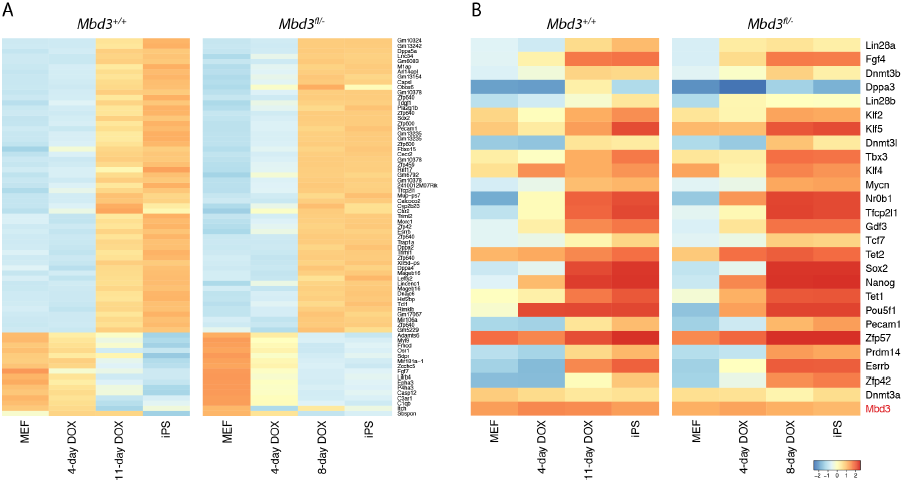
A) Microarray data from Rais *et al*. indicate similar gene expression patterns from *Mbd3*^+/+^ and *Mbd3*^fl/-^ cells during a timecourse of doxycycline-induced reprogramming (*Z*-score transformed log_2_ expression). B) Pluripotency factors show little variation between *Mbd3*^+/+^ and *Mbd3*^fl/-^ lines. *Mbd3* is expressed at comparable levels in both conditions (bottom row).

The greatest fold-change differences between experiment and control cells over the time series arise from a discordant set of genes devoid of canonical pluripotency regulators (Fig. 2A), suggesting the dominating effect to be due to biological variation expected from distinct and independently derived cell lines. However, the degree of such variability is impossible to assess in the absence of experimental replication.

**Figure 2.**
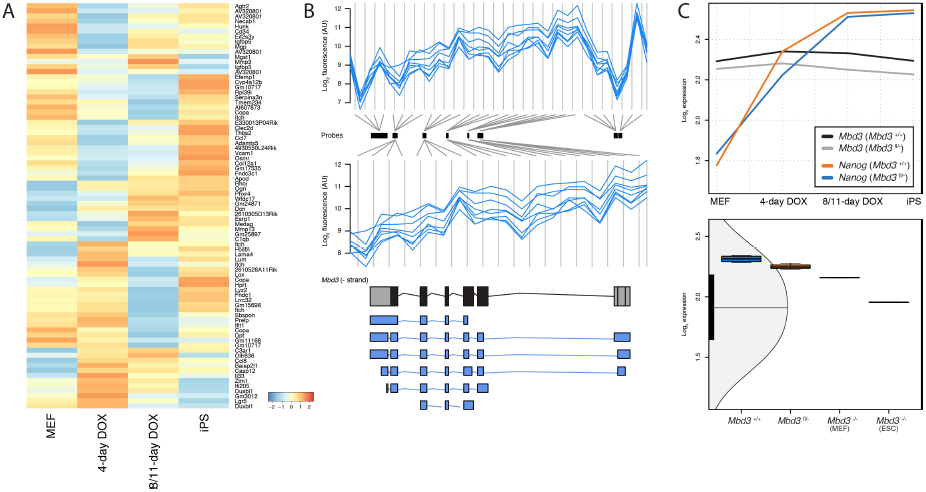
A) Genes exhibiting the most discrepant changes between *Mbd3*^+/+^ and *Mbd3*^fl/-^ cells profiled over the timecourse. B) Probe-level intensity data from the complete *Mbd3* probeset (upper data tracks) and reduced probeset excluding probes to exon 1 and UTRs remaining in the knockout allele (lower data tracks; see Methods). *Mbd3* locus and transcript isoforms are depicted below (antisense orientation). C) Comparable expression of *Mbd3* and *Nanog* in *Mbd3*^+/+^ and Mbd3^fl/-^ cells over the time series (top); *Mbd3* is transcribed in heterozygous and *Mbd3*^-/-^ MEFs at 85% and 66% wild-type levels, respectively (bottom). Continued expression in "null" cells may be due to reprogramming of MEF lines after Cre excision was performed, thereby selecting for iPS cell colonies where reversion to pluripotency was facilitated by the presence of an intact allele.

Rais *et al*. evaluated the potential of *Mbd3* depletion primarily in the *Mbd3*^fl/-^ heterozygous background without deleting the remaining allele. This was predicated on the notion that *Mbd3* displays hypomorphic expression, based on the authors’ estimate of protein abundance in *Mbd3*^fl/-^ cells at 20% that of wild-type levels. Markedly different results were obtained in two independent studies by our groups^9, 11^, where near wild-type Mbd3 protein abundance was measured from cells of this genotype regardless of the culture conditions used.

Analysis of the microarray data from Rais *et al*. shows *Mbd3* transcript levels in heterozygous cells to be 85% relative to *Mbd3*^+/+^ controls, consistent with the behavior previously observed. Although few experiments in Rais *et al*. involve cells in which the floxed allele had been deleted to assess reprogramming efficiency in null (*Mbd3*^-/-^) conditions, this was performed and expression data from those cells were included in the dataset. *Mbd3*^-/-^ MEFs profiled in the study express *Mbd3* transcript at 66% wild-type levels and 78% relative to *Mbd3*^fl/-^ cells (Fig. 2B,C), calling into question the effective depletion of Mbd3 protein and impairment of NuRD function as a causal factor contributing to the reported increase in reprogramming efficiency.

Much of the Rais *et al*. study makes use of a reporter of *Pou5f1* (*Oct4*) expression, consisting of the complete endogenous *Oct4* regulatory sequence linked to GFP^12^. Analysis of the ChIP-seq data provided by Rais *et al*. allows inspection of the promoter fragment used to regulate GFP expression in the reporter lines. The *Oct4* promoter region contains several well-characterized elements^13^, notably the proximal and distal enhancers (PE and DE). Their functions have been previously defined using <u>g</u>enomic *<u>O</u>ct4* <u>f</u>ragment (GOF)-18, an 18 kb intact sequence, and derivatives where regions encompassing each enhancer have been deleted (ΔPE and ΔDE)^14^.

Sequencing reads corresponding to the reporter transgene DNA map to the endogenous *Oct4* locus in the reference genome at high copy number (Fig. 3A). Alignments from *Mbd3*^fl/-^ cells are contiguous and span the entire promoter region. In contrast, a gap in read coverage is present in *Mbd3*^+/+^ cells corresponding to the segment deleted in the ΔPE construct (Fig. 3B). The intact GOF-18 construct is solely described in Rais *et al*. and indicated schematically in Extended Data Figure 3a (top). The full *Oct4* promoter is illustrated with PE and DE elements included, implying that all cells received this plasmid. In contrast, it is evident that *Mbd3*^fl/-^ and *Mbd3*^+/+^ control cells harbor different variants of GOF-18 reporter constructs.

**Figure 3.**
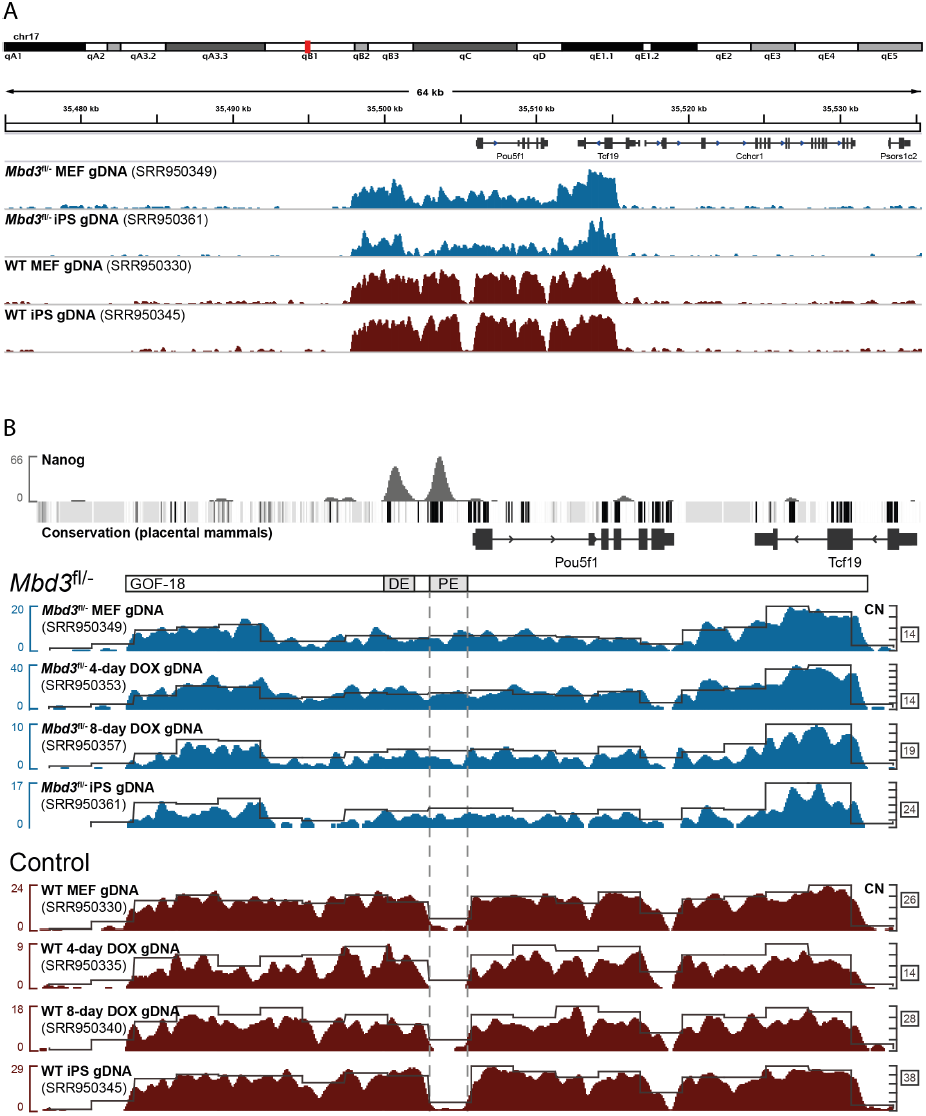
Sequencing data from whole-cell extract (WCE) genomic DNA libraries reveals numerous transgene copies relative to genomic background (A), with *Mbd3*^fl/-^ (blue) and *Mbd3*^+/+^ (red) *Oct4*-GFP reporter lines harboring intact GOF-18 and GOF-18 ΔPE constructs, respectively (B). Proximal and distal enhancer regions of the *Oct4* promoter are denoted, together with sequence conservation and Nanog binding site occupancy from an independent dataset^28^. Scales indicate read count (left) and transgene copy range estimated at 1 kb intervals (right).

The reprogramming system described in Rais *et al*. employed a polycistronic reprogramming cassette (STEMCCA)^15^ as well as a constitutive mCherry reporter used for quantification of colony sizes and single-cell deposition by flow sorting. To verify that GOF-18 APE *Mbd3*^+/+^ ChIP-seq data originated from the Oct4-GFP reporter cells used throughout the study, we identified sequencing reads corresponding to mCherry and parts of the STEMCCA construct design, including the internal ribosomal entry site (IRES) and 2A peptide sequences linking each reprogramming factor (Fig. 4).

**Figure 4.**
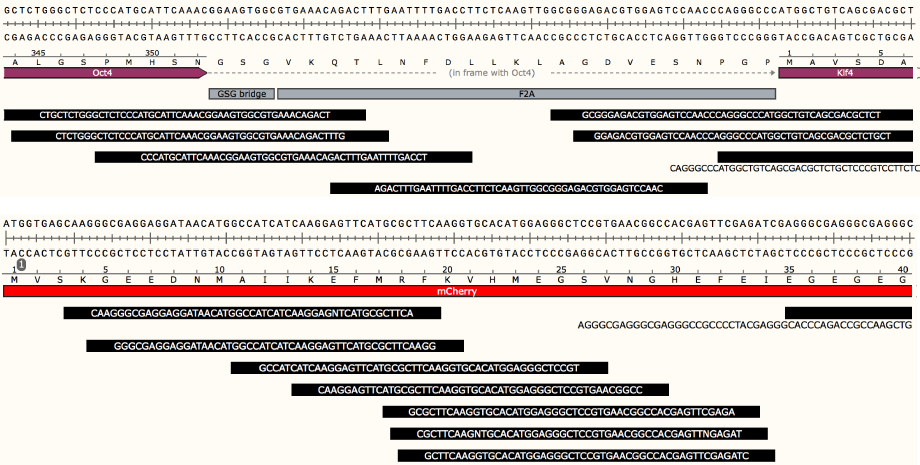
Sequencing data confirm *Mbd3*^+/+^ cells harbor the STEMCCA polycistronic cassette (top) and mCherry (bottom). Together with the GOF-18 ΔPE *Oct4*-GFP reporter (Fig. 3, 5) the complete reprogramming and reporter system is present, suggesting the use of these cells as controls in the assays listed in Table 1.

## Analysis and commentary

To properly evaluate the role of the NuRD complex by *Mbd3* depletion, both copies of the gene must be ablated. Rais *et al*. report hypomorphic expression from Mbd3^fl/-^ cells estimated at 20% wild-type levels, thereby justifying the use of a heterozygous cell line to represent a functional Mbd3 mutant. That assessment disagrees with our experience and the authors’ microarray data, where robust *Mbd3* expression is apparent in both *Mbd3*^fl/-^ and *Mbd3*^-/-^ lines. The latter finding may have arisen from incomplete Cre excision and/or positive selection of reprogramming-competent cells with an intact *Mbd3* allele. This suggests that differences in reprogramming kinetics are unlikely to be related to Mbd3 depletion, and indeed transcriptional states are comparable between the experiment and control cells profiled in the study.

Nonetheless, substantial improvements in reprogramming efficiency are described in Rais *et al*. Dramatic enhancement of pluripotency induction is reported from assays in which GFP was used as a readout for imaging and flow cytometry. Sequencing data from the study reveal that *Mbd3*^+/+^ cells were transfected with an *Oct4*-GFP reporter based on the GOF-18 ΔPE construct, whereas Mbd3^fl/-^ cells harbor an intact GOF-18 promoter fragment. *Oct4* is expressed in a wider repertoire of tissues and cell types than embryonic stem cells^16–19^ and reporters based on the intact GOF-18 construct display similarly broad activity^12, 14^. The PE is the most highly conserved region of the *Oct4* promoter in eutheria and also drives transcription in post-implantation embryos. Deleting the PE confines expression to naïve pluripotent cells, and thus a construct lacking the PE effectuates a much more stringent reporter of authentic reprogramming outcomes.

Differential application of a promiscuous test reporter and a considerably weaker control compromises the study design and undermines the conclusions drawn. An invalid experimental setup is imposed where no combination of *Oct4*-GFP reporter lines can be legitimately compared, as the two constructs have been applied in a mutually exclusive fashion to the experiment and control groups (Fig. 5). This applies to all ES-derived and iPS-derived MEFs where *Oct4*-GFP^+^ selection or quantification was used to establish differential reprogramming efficiency. No scientific motivation for comparative evaluation of alternate *Oct4*-GFP reporters is described in Rais *et al*., and use of the ΔPE variant is not declared. Thus the paper is lacking a key methodological disclosure essential for accurate interpretation of the results.

**Figure 5.**
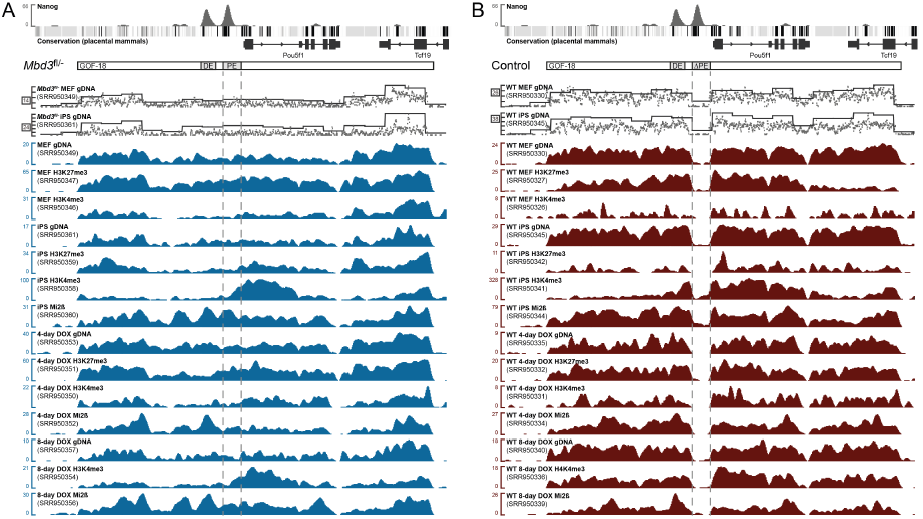
Data from equivalent ChIP-seq profiles indicate a 1:1 correspondence between the intact GOF-18 promoter construct applied to *Mbd3*^fl/-^ cells (left) versus the GOF-18 ΔPE variant present in *Mbd3*^+/+^ control cells (right).

The line of investigation presented in Rais *et al*. heavily relies on *Oct4-GFP* expression as a proxy for the reversion of somatic cells to pluripotency. Differences in reporter activity arising from the tandem use of intact GOF-18 and GOF-18 ΔPE constructs may have adversely affected a significant number of assays and conclusions presented in the study (Table 1). The trend depicted in Figure 2f of Rais *et al*. provides an illustrative example, where *Mbd3*^fl/-^ cells appear to revert to pluripotency at an accelerated rate relative to controls. Expression data from the study do not support that finding, which may have been construed on the basis of GFP output alone. *Mbd3*^+/+^ cells, where *Oct4*-GFP is driven by the ΔPE reporter, would be expected to yield profoundly reduced fluorescence signal relative to a variant based on the full promoter sequence.

**Table 1.** Exhibits from Rais *et al*. presenting data based on *Oct4*-GFP quantification to assess reprogramming efficiency.

| Figure | Assay | Interpretation |
| --- | --- | --- |
| 1f | Primordial germ cell derivation | p.66: "Single cell isolated *Mbd3*<sup>fl/-</sup> Oct4-GFP1 E8.5 PGCs from chimaeric mice were proficient in forming EG cell colonies and lines (.95% efficiency), whereas PGCs isolated from chimaeras that were generated by micro-injecting *Mbd3*<sup>+/+</sup> or *Mbd3*<sup>fl/-</sup> cells carrying an exogenous *Mbd3* transgene reprogrammed at less than 10% efficiency (Fig. 1f)." |
| Extended Data 3c | *Oct4*-GFP expression from reprogrammed cells and intermediates | p.66: "Notably, 95% of *Mbd3*<sup>fl/-</sup> and *Mbd3*<sup>-/-</sup> cells were Oct4-GFP<sup>+</sup> at day 10, whereas only levels up to 18% were observed in control *Mbd3*<sup>+/+</sup> fibroblasts (Fig. 2a)." |
| Extended Data 3d | *Oct4*-GFP expression from reprogrammed cells and intermediates |  |
| Extended Data 3f | *Oct4*-GFP expression from reprogrammed cells and intermediates |  |
| 2a | Reprogramming efficiency from *Mbd3*<sup>+/+</sup>, *Mbd3*<sup>fl/-</sup> and *Mbd3*<sup>-/-</sup> cells |  |
| 2b | Matrix of reprogramming outcomes (secondary iPS cells) | p.66: "Single cell sorting of secondary mCherry<sup>+</sup> *Mbd3*<sup>fl/-</sup> mouse embryonic fibroblasts (MEFs) and subsequent reprogramming in 2i/LIF plus doxycycline conditions reproducibly yielded 100% iPS cell derivation efficiency by day 8. Wild-type cells reprogrammed under identical conditions, no more than 20% of clones reactivated *Oct4*-GFP (Fig. 2b)." |
| Extended Data 3e | *Oct4*-GFP expression from reprogrammed cells and intermediates | p.67: "High single-cell reprogramming efficiency rates were obtained from a variety of adult progenitor and terminally differentiated cells (Fig. 2d and Extended Data Fig. 3e,f)." |
| 2e | Imaging of *Oct4*-GFP<sup>+</sup> vs mCherry<sup>+</sup> *Mbd3*<sup>+/+</sup> and *Mbd3*<sup>fl/-</sup> colonies | p.67: "By day 6 after doxycycline induction, .98% of *Mbd3*<sup>fl/-</sup> clonal populations reactivated the *Oct4*-GFP pluripotency marker, whereas only up to 20% efficiency was detected in control samples reprogrammed in identical growth conditions (Fig. 2e,f)." |
| 2f | Secondary reprogramming assay comparing *Oct4*-GFP<sup>+</sup> *Mbd3*<sup>+/+</sup> and *Mbd3*<sup>fl/-</sup> colonies (top) and cells (bottom) | p.67: "By day 6, approximately 85% of cells within each individual *Mbd3* clonal population became Oct4-GFP<sup>+</sup> cells, whereas <2% of cells within successfully reprogrammed *Mbd3*<sup>+/+</sup> clones turned on the *Oct4*-GFP marker (bottom panel in Fig. 2f)." |
| Extended Data 4a | Flow sorting of *Oct4*-GFP<sup>+</sup> *Mbd3*<sup>+/+</sup> and *Mbd3*<sup>fl/-</sup> cells | p.67: "Detection of Oct4-GFP by flow cytometry on polyclonal populations demonstrated similar iPS cell reprogramming kinetics (Extended Data Fig. 4a)." |
| Extended Data 4b | Time course of lentiviral transduction | p.67: "Re-infection with lentiviruses encoding *Mbd3*, but not *Mbd2*, before day 5 of reprogramming had a profound inhibitory effect on iPS cell generation from *Mbd3*<sup>fl/-</sup> MEFs, whereas re-infection after day 5 had a diminished effect (Extended Data Fig. 4b)." |
| 3b | Secondary reprogramming assay | p.67: "After the depletion of *Mbd3* expression, we were not able to isolate stable, partially reprogrammed cells that did not reactivate Oct4-GFP or Nanog-GFP and could be stably expanded in vitro, as typically can be obtained from OSKM-transduced wild-type somatic cells (Fig. 3b)." |
| 3c | Flow sorting of *Oct4*-GFP<sup>+</sup> cells | p.67: "Notably, by introducing *Mbd3* siRNA, all clones markedly turned on *Oct4*-GFP or *Nanog*-GFP pluripotency markers after continued OSKM expression in 2i/LIF (Fig. 3c)."<br>[only *Oct4*-GFP data are shown] |
| 5e | *Oct4*-GFP<sup>+</sup> cells in reprogramming assay | p.70: "Mbd3 mutants with a compromised ability to interact with OSKM reprogramming factors directly (Extended Data Fig. 9d) were deficient in reducing reprogramming efficiency of *Mbd3*<sup>fl/-</sup> somatic cells, supporting the notion that direct OSKM-Mbd3 interactions are important for inhibiting iPS cell formation (Fig. 5e)." |

During reprogramming, partially reverted intermediates are inherently produced en route to iPS cell colony formation. GFP expression from *Mbd3*^fl/-^ cells is unrestricted in these transitional states and is nonspecific for ground state pluripotency. This shortcoming is exacerbated by the authors’ use of serum replacement factors (e.g. KSR) in culture media, which abolishes specificity for naïve pluripotent cells conferred by inhibition of glycogen synthase kinase 3 (GSK3) and mitogen-activated protein kinase pathways^20^. Numerous *Oct4*-GFP transgene insertions present in *Mbd3*^fl/-^ (up to 24) and *Mbd3*^+/+^ (up to 38) cells were uncharacterized with respect to the regulatory context of integration sites, potentially leading to spurious GFP activation unrelated to complete reprogramming state or the expression of endogenous *Oct4*.

Appropriate controls were not implemented for the reprogramming system utilized in Rais *et al*. *Mbd3*^+/+^ cells did not constitute the parental line of the *Mbd3*^fl/-^ cells acquired for the study^2^; genetic modifications were delivered separately to *Mbd3*^fl/-^ and *Mbd3*^+/+^ primary and secondary donor cells; random transgene integrations were not assessed in any condition; and cell lines were independently derived. Reprogramming experiments were performed in permissive conditions and cells were transformed with excessive *Oct4*-GFP transgene copies such that fluorescence activation is likely to be misregulated. All of these factors contribute to considerable experimental variation and impair the determination of biological significance. Assays in which incompatible fluorescence reporters are directly compared cannot be considered valid.

Assessment of Mbd3/NuRD function in reprogramming must be conducted with validated *Mbd3*-null cells, compatible and equivalent genetic modifications in test and control conditions, rigorous evaluation of authentic pluripotent cells and reprogramming outcomes, and matched cell lines from an isogenic parental background. *Mbd3*^fl/-^ cells are not sufficient to assess the impact of *Mbd3* depletion, as cells of this genotype feature near wild-type transcript levels and protein abundance. In the absence of independent verification and in light of the deficiencies outlined above, results presented in Rais *et al*. describing 100% reprogramming efficiency based on the use of *Mbd3*^fl/-^ cells must be questioned as a potential artifact of the authors’ experimental system.

## Concluding remarks

We brought this matter to the attention of the authors, and upon receiving an unsatisfactory explanation for the disparities found, ultimately raised the issue with *Nature*. The editors declined to publish our exchange as a contribution to the Communications Arising section, and instead encouraged the authors to post a comment to the *Nature* website^21^. The comment makes readers aware of a difference in *Oct4*-GFP reporter usage, but the significance of this issue and its implications for the study as a whole are diminished. We therefore issue this letter as an expression of concern to investigators who would follow this work.

## Acknowledgements

We are grateful to Austin Smith and Wolfgang Huber for helpful discussions and advice.

## Methods

### Microarray data analysis

Affymetrix Mouse Gene 1.0 ST array data were obtained from GEO^22^ record GSE45352^8^ and processed with the *oligo* Bioconductor package^23^. Microarray data were normalized with the robust multi-array average (RMA) method^23^. Transcript clusters were mapped to mouse gene annotation based on release 78 of Ensembl^25^.

### *Mbd3* transcript expression

Microarray probesets targeting *Mbd3* were originally assigned a value of 1 in the `crosshyb_type` field of the Affymetrix design files, indicating each probe in the *Mbd3* transcript cluster (10370824) to be unique with respect to other putatively transcribed sequences targeted by the array. No additional perfect matches were found to any other mouse transcript annotated in Ensembl release 78, consistent with the assessment of cross-hybridization potential carried out at design time. Heterozygous knockout (*Mbd3*^fl/-^) cells had been targeted such that exons 2–7 were replaced with the β-geo selection marker, leaving exon 1 and UTR sequences intact^10^. To discount residual contribution from the non-functional allele, sense-orientation probe sequences were mapped to the reverse complement of *Mbd3* genomic DNA, and probes corresponding to exon 1 (84510, 233909, 995596, 1042262), 5′ UTR (314091, 646154, 26469) and 3′ UTR (1028146, 391255, 585086, 333495) were deleted from the pd.mogene.1.0.st.v1 annotation database^26^ prior to normalization. Expression levels were estimated as described above from the remaining 20 of 31 original probes spanning the *Mbd3* locus. Probe-level data were plotted with the *GenomeGraphs* Bioconductor package^27^.

### ChIP-seq data analysis

Illumina sequencing data deposited under accessions SRP028718^8^ and SRX000545^28^ were obtained from the Sequence Read Archive^29^ and aligned to the mouse genome GRCm38 (mm10) using BWA^30^, allowing permissive treatment of low-quality base calls (`–l 2 5 –q 20`). For conservative copy number estimation, duplicate reads likely arising from PCR amplification were removed with Picard^31^, and suboptimal alignments (`–q 10`) filtered with SAMtools^32^. Focal gains corresponding to transgene insertions were estimated from genomic DNA (WCE, whole-cell extract) samples, accounting for G/C content^33^ and assuming ploidy = 2 over windows of 1–10 kb. Read density was computed with F-Seq^34^ and visualized in the Integrative Genomics Viewer^35^.

## References

1. Zhang, Y., et al. Analysis of the NuRD subunits reveals a histone deacetylase core complex and a connection with DNA methylation. Genes Dev. 13, 1924–35 (1999)

2. Kaji, K., et al. The NuRD component Mbd3 is required for pluripotency of embryonic stem cells. Nat Cell Biol. 8, 285–92 (2006)

3. Reynolds, N., et al. NuRD-mediated deacetylation of H3K27 facilitates recruitment of Polycomb Repressive Complex 2 to direct gene repression. EMBO J. 31, 593–605 (2012)

4. Costa, Y., et al. NANOG-dependent function of TET1 and TET2 in establishment of pluripotency. Nature 495, 370–4 (2013)

5. Gagliardi, A., et al. A direct physical interaction between Nanog and Sox2 regulates embryonic stem cell self-renewal. EMBO J. 32, 2231–47 (2013)

6. van den Berg, D.L., et al. An Oct4-centered protein interaction network in embryonic stem cells. Cell Stem Cell 6, 369–81 (2010)

7. Pardo, M., et al. An expanded Oct4 interaction network: implications for stem cell biology, development, and disease. Cell Stem Cell 6, 382–95 (2010)

8. Rais, Y. et al. Deterministic direct reprogramming of somatic cells to pluripotency. Nature 502, 65–70 (2013)

9. dos Santos, R.L., et al. MBD3/NuRD facilitates induction of pluripotency in a context-dependent manner. Cell Stem Cell 15, 102–10 (2014)

10. Hendrich, B., Guy, J., Ramsahoye, B., Wilson, V.A. & Bird, A. Closely related proteins MBD2 and MBD3 play distinctive but interacting roles in mouse development. Genes Dev. 15, 710–23 (2001)

11. Reynolds, N., et al. NuRD suppresses pluripotency gene expression to promote transcriptional heterogeneity and lineage commitment. Cell Stem Cell 10, 583–94 (2012)

12. Kirchhof, N., et al. Expression pattern of Oct-4 in preimplantation embryos of different species. Biol Reprod. 63, 1698–705 (2000)

13. Sylvester, I. & Scholer, H.R. Regulation of the Oct-4 gene by nuclear receptors. Nucleic Acids Res. 22, 901–11 (1994)

14. Yeom, Y.I., et al. Germline regulatory element of Oct-4 specific for the totipotent cycle of embryonal cells. Development 122, 881–94 (1996)

15. Sommer, C.A., et al. Induced pluripotent stem cell generation using a single lentiviral stem cell cassette. Stem Cells 27, 543–9 (2009)

16. Kehler, J., et al. Oct4 is required for primordial germ cell survival. EMBO Rep. 5, 1078–83 (2004)

17. Han, D.W., et al. Epiblast stem cell subpopulations represent mouse embryos of distinct pregastrulation stages. Cell 143, 617–27 (2010)

18. Frum, T., et al. Oct4 cell-autonomously promotes primitive endoderm development in the mouse blastocyst. Dev Cell 25, 610–22 (2013)

19. Le Bin, G.C., et al. Oct4 is required for lineage priming in the developing inner cell mass of the mouse blastocyst. Development 141, 1001–10 (2014)

20. Silva, J., et al. Promotion of reprogramming to ground state pluripotency by signal inhibition. PLoS Biol. 6, e253 (2008)

21. http://www.nature.com/nature/journal/v502/n7469/full/nature12587.html#comment-64027

22. Barrett, T., et al. NCBI GEO: archive for functional genomics data sets-update. Nucleic Acids Res. 41, D991-5 (2013)

23. Carvalho, B.S. & Irizarry, R.A. A framework for oligonucleotide microarray preprocessing. Bioinformatics 26, 2363–7 (2010)

24. Irizarry, R.A., et al. Exploration, normalization, and summaries of high density oligonucleotide array probe level data. Biostatistics 4, 249–264 (2003)

25. Cunningham, F., et al. Ensembl 2015. Nucleic Acids Res. gku1010 (2014)

26. Carvalho, B. pd.mogene.1.0.st.v1: platform design info for Affymetrix MoGene-1_0-st-v1. R package version 3.10.0.

27. Durinck, S., Bullard, J., Spellman, P.T. & Dudoit, S. GenomeGraphs: integrated genomic data visualization with R. BMC Bioinformatics 10, 2 (2009)

28. Chen, X., et al. Integration of external signaling pathways with the core transcriptional network in embryonic stem cells. Cell 133, 1106–17 (2008)

29. Leinonen, R., Sugawara, H., Shumway, M; International Nucleotide Sequence Database Collaboration. The sequence read archive. Nucleic Acids Res. 39, D19–21 (2011)

30. Li, H. & Durbin, R. Fast and accurate short read alignment with Burrows-Wheeler Transform. Bioinformatics 25, 1754–60 (2009)

31. http://broadinstitute.github.io/picard

32. Li, H., et al. The Sequence Alignment/Map (SAM) format and SAMtools. Bioinformatics 25, 2078–9 (2009)

33. Boeva, V., et al. Control-free calling of copy number alterations in deep-sequencing data using GC-content normalization. Bioinformatics 15, 268–9 (2011)

34. Boyle, A.P., Guinney, J., Crawford, G.E. & Furey, T.S. F-Seq: a feature density estimator for high-throughput sequence tags. Bioinformatics 24, 2537–8 (2008)

35. Thorvaldsdóttir, H., Robinson, J.T. & Mesirov, J.P. Integrative Genomics Viewer (IGV): high-performance genomics data visualization and exploration. Brief Bioinform. 14, 178–92 (2012)

